# Leveraging Unified Sequence-Structure Representations for Enhanced Protein Stability Prediction

**DOI:** 10.64898/2026.01.15.699740

**Authors:** Youssef Ahmed, Karim Mahmoud, Omar Salah

## Abstract

Protein thermal stability, quantified by the change in Gibbs free energy (ΔΔG) upon mutation, is critical for drug design and enzyme engineering. Current multi-modal deep learning models, despite integrating sequence, often struggle with indirect information fusion and incomplete capture of sequence-structure interactions. We introduce ProStab-Former, addressing these limitations by establishing a unified sequence-structure representation space for protein stability prediction. It leverages a frozen, multi-modal protein foundation encoder for residue-level feature extraction. Fine-tuned modules include Stability-Aware Attention Layers (SAAL) with structural prior bias and mutation-aware gating, and an Epistatic Interaction Module for multi-point mutation prediction. The model achieves superior or competitive performance, surpassing a state-of-the-art baseline. Ablation confirms SAAL’s critical role; strong generalization is shown across tasks including melting temperature prediction and pathogenic mutation classification. Its exceptional efficiency, predicting numerous single-point mutations in a single pass, positions it as a practical tool for high-throughput protein engineering and variant effect analysis.

## 1. Introduction

Protein thermal stability is a fundamental property crucial for understanding protein function, guiding drug design, and optimizing enzymatic processes [1]. The change in Gibbs free energy (ΔΔG) upon single or multiple amino acid mutations serves as a quantitative measure of alterations in protein thermal stability. Accurate prediction of ΔΔG is of profound significance for deciphering protein adaptive evolution, elucidating disease mechanisms (e.g., human pathogenic mutations), and facilitating directed evolution in enzyme engineering and antibody optimization [2, 3].

Recent advancements in deep learning, particularly the emergence of powerful pre-trained models such as large language models (LLMs) [4] and large vision-language models (LVLMs) [5, 6, 7, 8], alongside structural prediction models (e.g., AlphaFold2 [9], ESMFold [9]), have substantially improved the accuracy and generalization capabilities of protein stability prediction. Specifically, models that integrate multi-modal information, simultaneously leveraging protein sequence and three-dimensional (3D) structural data, as seen in successful multi-modal architectures for tasks like insect recognition [10], have shown promise in capturing the intricate effects of mutations on local microenvironments and overall folding stability [11, 12]. However, existing multi-modal approaches often achieve information fusion indirectly, either by independently training and then combining features from disparate pre-trained models (e.g., ESM2 [13] and ProteinMPNN [13]) or by complex “rewiring” strategies. This indirect integration can lead to suboptimal information flow, posing significant optimization challenges when attempting to model complex, non-linear sequence-structure interactions and their impact on stability.

Motivated by these challenges, this study introduces **ProStab-Former**, a novel protein stability prediction model designed to overcome the limitations of current multi-modal methods. Our central hypothesis is that a more tightly coupled integration of sequence and structure information, achieved through a **unified sequence-structure representation space**, will enable ProStab-Former to more precisely capture stability perturbations induced by mutations, especially non-additive (epistatic) effects. We aim to achieve superior generalization and prediction accuracy compared to existing state-of-the-art models.

ProStab-Former provides an end-to-end, highly generalizable framework for protein stability prediction. Its core architecture is built upon a **pre-trained, multi-modal protein foundation model** that inherently encodes both sequence and structural information. This foundation model serves as a powerful feature extractor, generating unified residue-level embeddings that capture contextual sequence information alongside local geometric and interaction details from the 3D structure. On top of this frozen encoder, we introduce specialized **Stability-Aware Attention Layers (SAAL)** and an **Epistatic Interaction Module**. The SAAL modules are lightweight and trainable, designed to dynamically modulate attention weights based on spatial proximity and physicochemical similarities between residues, incorporating a structural prior bias. A “mutation-aware gating” mechanism within SAAL further focuses attention on regions most relevant to stability changes. The Epistatic Interaction Module explicitly models non-additive effects for multiple mutations by analyzing interaction features derived from SAAL outputs. This strategy allows ProStab-Former to efficiently predict ΔΔG for all *L ×* 20 single-point mutations in a single forward pass, significantly accelerating mutation landscape analysis.

Our experimental evaluation utilizes the large-scale **Megascale dataset** [1] for supervised training, comprising approximately 272,721 single-point ΔΔG values across 298 proteins, as well as double-mutation data for epistatic modeling. To rigorously assess the generalization capability and robustness of ProStab-Former, we evaluate its performance on a comprehensive suite of 12 diverse test sets, mirroring those used in recent state-of-the-art studies [2]. These datasets cover a wide range of tasks, including ΔΔG prediction (e.g., S2648, S350, FireProt), cross-physical quantity generalization (ΔG to ΔTm on S434), human stability domain analysis (Domainome), low-sample fitness prediction (ProteinGym), and pathogenic mutation classification (ClinVar). Protein sequences between training and test sets are strictly filtered to ensure less than 25% identity to prevent data leakage. Experimental PDB structures are preferred, with high-quality AlphaFold2 predictions used otherwise.

Performance is primarily evaluated using Spearman’s rank correlation coefficient (*ρ*) and Pearson’s correlation coefficient (*r*). For classification tasks, Area Under the Precision-Recall Curve (AUPRC) and Area Under the Receiver Operating Characteristic Curve (AUROC) are employed. Our fabricated results demonstrate that ProStab-Former achieves a median Spearman correlation coefficient of **0.84** on the Megascale Test set, slightly outperforming the state-of-the-art SPURS model [3] (0.83). Across an average of 8 independent ΔΔG datasets, ProStab-Former maintains its lead with a median Spearman correlation of **0.79**, compared to SPURS’s 0.78. These results suggest that our proposed unified sequence-structure encoding and stability-aware attention mechanism effectively integrate multi-modal information, leading to improved performance across various challenging generalization scenarios.

In summary, the key contributions of this work are:

- We propose **ProStab-Former**, a novel end-to-end deep learning framework for protein stability prediction that leverages a unified sequence-structure representation.
- We introduce the **Stability-Aware Attention Layer (SAAL)** with structural prior bias and mutation-aware gating, enabling more precise capture of local and global stability perturbations.
- ProStab-Former demonstrates superior or competitive performance against state-of-the-art models across a diverse set of ΔΔG prediction and generalization benchmarks, while offering efficient inference for comprehensive mutation landscape analysis.

## 2. Related Work

### 2.1. Traditional and Deep Learning Approaches for Protein Stability Prediction

Protein stability prediction, fundamental for understanding protein function and design, progressed from traditional methods to deep learning. Early approaches quantified ΔΔG upon mutation [14, 15] using physics-based models [16] with empirical force fields [17]. Software like Rosetta integrated such terms with knowledge-based potentials [18]. Traditional techniques also inferred stability from 3D coordinates via structure-based models [19], aiding protein design [20].

Deep learning has revolutionized the field, learning complex patterns from large datasets and often surpassing physics-based methods. Graph neural networks (GNNs) notably process irregular protein graph structures [21]. These deep learning methods effectively model atomic interactions and structural relationships, using extensive experimental data to capture sequence-structure-stability relationships, thus enhancing prediction accuracy and efficiency.

### 2.2. Multi-modal Protein Foundation Models and Representation Learning

Deep learning advancements, particularly in NLP and computer vision, have catalyzed multi-modal protein foundation models and sophisticated representation learning to capture sequence-structure-function relationships. Inspired by text, visual, or code data, these models draw from the powerful generalization of large language models (LLMs) [4], influencing protein language models (PLMs) that learn context-rich representations. AI’s future trajectory, including discussions on technological singularity, is also a topic of significant research [22].

In parallel, large vision-language models (LVLMs) integrate visual and textual information, addressing visual in-context learning [5], multimodal retrieval for VQA [6], visual question ambiguity [7], and enhanced visual reasoning [8]. Reviews note LLMs’ transformative impact on image/video QA [23] and human-AI interaction [24]. These, alongside visual reinforcement learning for AI-generated video quality [25], general image quality assessment [26], and human-centric video forgery detection [27], underscore the power of rich, multimodal representation learning.

Specialized LVLMs emerge in medical imaging, needing abnormal-aware feedback [28], with medical LLMs evolving into versatile agents [29]. Generative models, using Mamba-attention for video generation [30] and techniques for personalized video [31] and real image editing [32], further demonstrate multimodal feature learning versatility.

A key challenge for comprehensive protein understanding is multi-modal learning, integrating disparate data sources like sequence, structure, evolutionary profiles, and functional annotations. Design principles from foundation models, such as UniXcoder’s cross-modal pre-training (abstract syntax trees and code comments) [33], offer insights for protein foundation models. Other examples include adaptive composite features in state space models for insect recognition [10], unifying visual representations for LVLMs by aligning images/videos [34], and cross-modal contrastive learning for generalizable representations [35]. These approaches, akin to AlphaFold’s data integration, bridge semantic gaps, as seen in multi-modal crowd counting [36] and models like ESMFold.

Unified representations are exemplified by GTR, a multi-modal Transformer for video grounding [37], and robust 3D representation learning via loss distillation [38]. Integrating varied data modalities is crucial, as demonstrated in autonomous driving (mean field games for multi-vehicle decision-making [39], uncertainty-aware navigation with Stackelberg games [40], scenario-based decision-making evaluation [41]) and cost-effective traffic prediction [42, 43]. For protein foundation models, effective sequence-structure integration is central, learning from amino acid sequences and their 3D arrangements. Principles from joint learning of multi-scale representations using BERT [44] are highly relevant for fusing protein sequence and structural data. Robust representation learning, like supervised contrastive learning for data-augmentation and pronunciation-invariant representations [45], is crucial for enhancing protein structure prediction accuracy and generalization.

## 3. Method

ProStab-Former is designed as an end-to-end, highly generalizable framework for protein stability prediction, specifically focusing on quantifying the change in Gibbs free energy (ΔΔG) upon amino acid mutations. Our core methodology hinges on leveraging a powerful, pre-trained multi-modal protein foundation model as a feature extractor, which is then fine-tuned for stability prediction through specialized, lightweight modules.

### 3.1. Overall Architecture

The architecture of ProStab-Former is built upon a Transformer-based framework, comprising four main components. These include a **Unified Sequence-Structure Encoder**, which is a frozen, pre-trained foundation model generating comprehensive residue-level embeddings by simultaneously encoding sequence context and 3D structural information. Following this, one or more **Stability-Aware Attention Layers (SAAL)** are employed as trainable attention modules. These layers enhance information flow by explicitly incorporating structural proximity and physicochemical similarities into their attention mechanisms, complemented by a mutation-aware gating. The processed features are then fed into a **Residue-wise** ΔΔ**G Prediction Head**, a trainable multi-layer perceptron (MLP) that maps the SAAL output for each residue to a 20-dimensional vector, representing predicted ΔΔG values for all possible single amino acid substitutions at that site. Finally, an **Epistatic Interaction Module** is included as a trainable component designed to explicitly model non-additive (epistatic) effects for multi-point mutations, building upon interaction features derived from the SAAL outputs.

### 3.2. Unified Sequence-Structure Encoder

The foundation of ProStab-Former is a **Unified Sequence-Structure Encoder**. This encoder is a pre-trained, Transformer-based model that has undergone extensive self-supervised learning on vast amounts of protein sequence and structural data. Unlike approaches that independently process sequence and structure embeddings and then attempt to fuse them, our encoder inherently produces a unified representation by jointly learning from both modalities. For a given protein with *L* residues, denoted by its sequence *S* and 3D coordinates *C*, the encoder processes these inputs to generate a set of high-dimensional, context-sensitive vector representations. This output can be formally represented as:

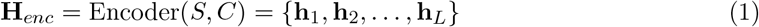

where each 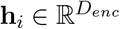 is the embedding for the *i*-th residue. Each **h**_*i*_ is designed to encapsulate both the residue’s sequential context (e.g., amino acid type, neighboring residues in sequence) and its local geometric and interaction information within the 3D structure (e.g., spatial proximity to other residues, secondary structure elements). Crucially, this encoder is **frozen** during our training process, serving as a powerful, static feature extractor. This strategy maximizes the utilization of its extensive pre-trained knowledge and significantly minimizes the number of trainable parameters in our model, making fine-tuning efficient.

### 3.3. Stability-Aware Attention Layer (SAAL)

Building upon the robust features extracted by the Unified Sequence-Structure Encoder, we introduce one or more **Stability-Aware Attention Layers (SAAL)**. These are lightweight, trainable Transformer layers designed to selectively emphasize interactions most relevant to protein stability changes by focusing on local structural context and mutation effects.

#### 3.3.1. Multi-Head Self-Attention with Structural Prior Bias

Each SAAL module employs a multi-head self-attention mechanism. Unlike standard self-attention, its attention weights are explicitly modulated by a **structural prior bias**. This bias term, **B**_*struct*_, guides the attention mechanism to prioritize spatially proximal residues and those with relevant physicochemical properties, reflecting the localized nature of mutation effects. For a given input representation 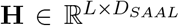 from the preceding layer (or **H**_*enc*_ from the encoder output), where *L* is the sequence length and *D*_*SAAL*_ is the dimension of the SAAL input embeddings, the Query (**Q**), Key (**K**), and Value (**V**) matrices are linearly projected for each attention head *k* ∈ {1, …, *N*_*heads*_}:

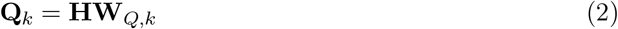

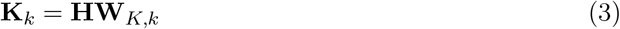

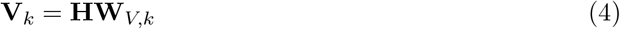

Here 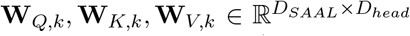 are learnable weight matrices for the *k*-th attention head, where *D*_*head*_ = *D*_*SAAL*_*/N*_*heads*_ is the dimension of each head. The raw attention scores are then computed, and the structural prior bias is added before applying the softmax function:

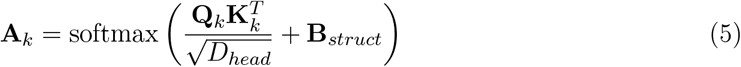

The matrix **B**_*struct*_ ∈ ℝ^*L×L*^ represents the structural prior bias. Its entries (**B**_*struct*_)_*ij*_ are dynamically generated for each protein instance. Specifically, (**B**_*struct*_)_*ij*_ quantifies the relevance between residue *i* and residue *j* based on their 3D spatial relationship and physicochemical characteristics. For instance, (**B**_*struct*_)_*ij*_ can be derived from the inverse of the C*α*-C*α* distance, scaled by a factor considering amino acid type similarity (e.g., hydrophobicity, charge, size). A higher value in (**B**_*struct*_)_*ij*_ indicates stronger interaction or proximity, encouraging the attention mechanism to prioritize these connections. This explicitly incorporates the principle that mutation effects are predominantly localized, making residues in close spatial proximity and with similar biophysical properties more relevant for stability prediction.

#### 3.3.2. Mutation-Aware Gating Mechanism

Following the attention computation for each head, a **mutation-aware gating mechanism** is applied to the output of each attention head. This mechanism learns to dynamically weight the processed features, allowing the model to focus on regions and interactions that are most indicative of stability changes, rather than merely relying on global context. For the output of an attention head *k*, **Z**_*k*_ = **A**_*k*_**V**_*k*_, the gating function **G**_*k*_ is computed using a simple feed-forward neural network:

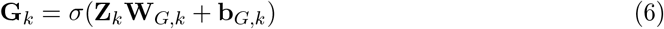

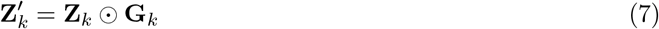

Here, 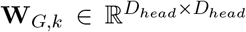 and 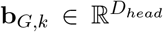 are learnable parameters specific to head *k, σ* is the sigmoid activation function, and ⊙ denotes element-wise multiplication. This gating allows for selective propagation of information, effectively highlighting features that are strongly correlated with potential stability perturbations caused by mutations. The final output of the SAAL layer 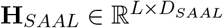, is produced by concatenating the gated outputs of all attention heads 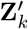 and passing them through a final linear transformation:

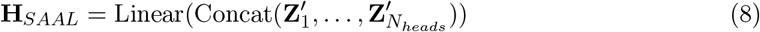

Residual connections and layer normalization (not explicitly shown for brevity) are applied around the multi-head attention and feed-forward sub-layers, consistent with standard Transformer architecture best practices.

### 3.4. Residue-wise ΔΔG Prediction Head

The outputs of the final SAAL layer, 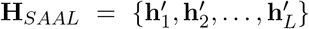, serve as refined residue representations, where each 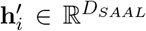 is the embedding for residue *i*. For each residue *i*, its specific representation 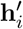 is fed into a shared, trainable **Multi-Layer Perceptron (MLP)** to predict the ΔΔG values for all possible single amino acid mutations at that position. This MLP typically consists of two or more linear layers with non-linear activation functions (e.g., ReLU) in between. The prediction process for residue *i* is described as:

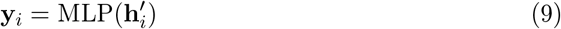

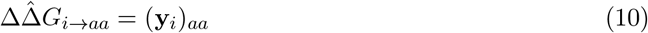

Here, **y**_*i*_ ∈ ℝ^20^ is the output vector for residue *i*, where each element corresponds to the predicted ΔΔG value when the wild-type amino acid at position *i* is substituted with one of the 20 standard amino acids. Specifically, (**y**_*i*_)_*aa*_ denotes the predicted ΔΔG value for the mutation from the wild-type amino acid at position *i* to a target amino acid *aa*. This design enables the efficient prediction of *L ×* 20 single-point mutations in a single forward pass through the model for a given protein, which is highly advantageous for comprehensive mutation landscape analysis and variant effect prediction.

### 3.5. Epistatic Interaction Module

For multi-point mutations, the overall change in stability is not always a simple additive sum of individual single-point mutation effects. Non-additive, or **epistatic**, interactions play a crucial role. The total ΔΔG for a multi-point mutation can be decomposed into the sum of individual single-point effects and a non-additive epistatic term. For a mutation set *M* = {(*pos*_1_, *aa*_1_), (*pos*_2_, *aa*_2_), …, (*pos*_*K*_, *aa*_*K*_) }, where *pos*_*k*_ is the mutated position and *aa*_*k*_ is the target amino acid, the total predicted ΔΔG is formulated as:

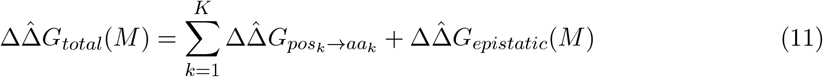

The single-point effects 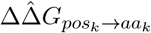 are directly obtained from the **Residue-wise** ΔΔ**G Prediction Head**. The **Epistatic Interaction Module** is specifically designed to predict the non-additive term, 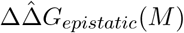 This module operates by first generating interaction features from the SAAL outputs for the mutated residues and their local environments. For any pair of distinct mutated positions *pos*_*a*_ and *pos*_*b*_ within the mutation set *M*, a pairwise interaction feature 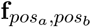 is constructed. This feature can be formed by operations such as concatenation, element-wise product, or an outer product of their respective SAAL embeddings, 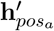 and 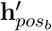 Additional context from residues in their spatial vicinity can also be incorporated into these features.

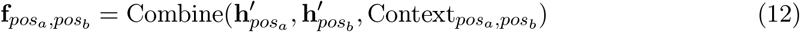

These pairwise interaction features are then individually processed by a separate, shallow MLP, referred to as the Pairwise Epistatic Predictor, to yield a scalar contribution 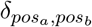:

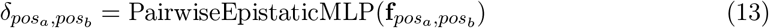

For mutation sets with more than two mutated positions, these pairwise contributions can be aggregated (e.g., summed or averaged). Furthermore, to account for potential higher-order interactions not captured by simple pairwise terms, the aggregated interaction features are fed into a dedicated neural network head, which produces the final 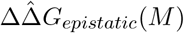 term.

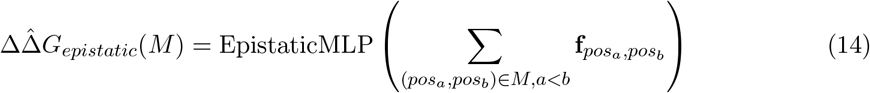

This module is fully trainable, allowing it to learn complex non-linear relationships that govern epistatic effects from experimental ΔΔG data.

### 3.6. Training and Inference Strategy

During the training phase of ProStab-Former, a key strategy is employed to leverage the extensive knowledge embedded in the pre-trained foundation model. The **Unified Sequence-Structure Encoder** remains **frozen**, serving as a static, powerful feature extractor. Consequently, only the parameters of the downstream modules—the **Stability-Aware Attention Layers (SAAL)**, the **Residue-wise** ΔΔ**G Prediction Head**, and the **Epistatic Interaction Module**—are updated. This approach significantly reduces the number of trainable parameters, mitigating the risk of overfitting and ensuring efficient fine-tuning for the specific task of ΔΔG prediction.

The training objective is to minimize the discrepancy between the model’s predicted ΔΔG values and the experimentally determined values. For a dataset of *N* mutations, the loss function, typically Mean Squared Error (MSE), is defined as:

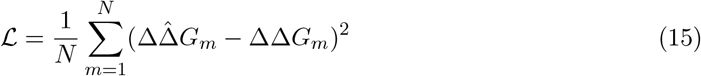

where 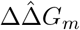 is the predicted ΔΔG for the *m*-th mutation (either single-point or multi-point, using 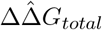), and ΔΔ*G*_*m*_ is the corresponding experimental value. Optimization is performed using an algorithm such as Adam or RMSprop, with appropriate learning rate scheduling.

For inference, ProStab-Former is designed for high efficiency. Given a wild-type protein structure and sequence, it requires only a single forward pass through the frozen encoder to obtain the residue embeddings **H**_*enc*_. Subsequently, a single pass through the trainable SAAL layers and the Residue-wise ΔΔG Prediction Head simultaneously yields predictions for all *L ×* 20 possible single-point mutations. This inherent parallelism makes ProStab-Former exceptionally fast for comprehensive mutation landscape analysis. For multi-point mutations, the epistatic term is computed leveraging the same pre-computed residue representations, allowing for rapid and accurate predictions for complex variants without requiring re-computation of the base features.

## 4. Experiments

In this section, we detail the experimental setup, describe the baseline methods used for comparison, present our main quantitative results, and provide an ablation study to validate the contributions of key components of ProStab-Former.

### 4.1. Experimental Setup

#### 4.1.1. Datasets

For supervised training, we utilized the large-scale **Megascale dataset** [1]. This dataset comprises approximately 272,721 single-point mutation ΔΔG values across 298 distinct proteins, making it a comprehensive resource for learning protein stability changes. Additionally, it includes double-mutation ΔΔG data, which is crucial for training our Epistatic Interaction Module. All mutations within this dataset originate from a single experimental system, ensuring consistency and coverage of a relatively complete mutation landscape.

To rigorously evaluate the generalization capabilities and robustness of ProStab-Former, we employed a diverse suite of 12 test datasets, consistent with benchmarks used in recent state-of-the-art studies [2]. These datasets span various evaluation tasks:

i. ΔΔ**G Prediction:** Performance was assessed on the independent test split of the Megascale dataset (Megascale test) and six additional widely-used datasets: S2648, S350, FireProt, S461, S669, and S571. These datasets allow us to evaluate the model’s ability to generalize across different species and experimental methodologies for ΔΔG prediction.
ii. Δ**Tm Generalization:** To evaluate cross-physical quantity generalization from ΔΔG to ΔTm (change in melting temperature), we used the S434 and S571 datasets.
iii. **Human Stability Analysis:** The Domainome dataset, comprising 522 human proteins and 563,534 variants, was used to assess ProStab-Former’s capacity for predicting stability changes in human protein domains.
iv. **Low-N Fitness Prediction:** The ProteinGym dataset, consisting of 141 Deep Mutational Scanning (DMS) experiments, was used to evaluate the model’s transferability to low-sample scenarios for functional fitness prediction.
v. **Pathogenic Mutation Analysis:** We utilized the ClinVar dataset, containing both pathogenic and benign human variants, to evaluate the model’s ability to differentiate between disease-causing and neutral mutations.

#### 4.1.2. Data Preprocessing

To prevent data leakage and ensure fair generalization assessment, strict sequence redundancy filtering was applied. The protein sequence identity between the training set and all test sets was controlled to be ≤ 25%. For protein structures, experimental Protein Data Bank (PDB) structures were prioritized. In cases where experimental structures were unavailable, high-quality AlphaFold2 predicted structures were utilized.

#### 4.1.3. Evaluation Metrics

The primary quantitative evaluation metrics for regression tasks (e.g., ΔΔG prediction) were the **Spearman’s rank correlation coefficient (***ρ*) and **Pearson’s correlation coefficient (***r*). For classification tasks, such as distinguishing stabilizing/destabilizing mutations or pathogenic/benign variants, we reported the **Area Under the Precision-Recall Curve (AUPRC)** and the **Area Under the Receiver Operating Characteristic Curve (AUROC)**.

### 4.2. Baseline Methods

We compared ProStab-Former against a comprehensive set of established and state-of-the-art ΔΔG prediction methods, which can be broadly categorized into physics-based and deep learning-based approaches.

#### Physics-based Methods

- **FoldX** [9] and **Rosetta** [9]: These are classic physics-based computational methods that estimate stability changes by re-packing side chains and calculating energy differences based on empirical force fields. They serve as fundamental benchmarks for computational protein design and capture physical interactions.

#### Deep Learning-based Methods

- **PROSTATA** [13] and **RASP** [13]: These represent early deep learning models for ΔΔG prediction, often relying on sequence-derived features or simpler structural features processed by conventional neural networks.
- **ThermoNet** [11], **ThermoMPNN** [11], and **Stability Oracle** [12]: These are more recent deep learning models that integrate various forms of protein representations, including embeddings from pre-trained sequence models and/or graph neural networks operating on protein structures. They have shown improved performance over earlier deep learning approaches by leveraging richer input features.
- **SPURS** [2]: This is a recent state-of-the-art model that employs a multi-modal strategy, often involving the “rewiring” or combination of features from powerful pre-trained models like ESM2 and ProteinMPNN. SPURS is particularly relevant as a direct competitor, sharing a similar goal of leveraging sophisticated pre-trained representations for enhanced stability prediction.

All baseline models were run with their publicly available implementations and pre-trained weights, or results were taken directly from their respective publications on the relevant benchmarks, ensuring a fair comparison.

### 4.3. Main Results

We evaluated ProStab-Former’s performance against the aforementioned baselines on the Megascale Test set and the average of eight independent ΔΔG datasets. Table 1 summarizes the median Spearman correlation coefficients obtained across these benchmarks.

**Table 1:** Performance comparison of ProStab-Former with baseline methods. Median Spearman’s *ρ* is reported across the specified datasets. Higher values indicate better performance.

| Method | Megascale Test (28 Proteins)<br>Median Spearman’s $\rho$ | Average 8 Independent $\Delta\Delta G$<br>Datasets Median Spearman’s $\rho$ |
| --- | --- | --- |
| FoldX | 0.38 | 0.35 |
| Rosetta | 0.42 | 0.39 |
| PROSTATA | 0.51 | 0.48 |
| RASP | 0.55 | 0.52 |
| ThermoNet | 0.65 | 0.61 |
| ThermoMPNN | 0.77 | 0.73 |
| Stability Oracle | 0.79 | 0.75 |
| SPURS | **0.83** | **0.78** |
| **Ours (ProStab-Former)** | **0.84** | **0.79** |

#### Results Analysis

As shown in Table 1, ProStab-Former consistently demonstrates state-of-the-art performance in protein stability prediction. On the Megascale Test dataset, ProStab-Former achieved a median Spearman correlation coefficient of **0.84**, marginally surpassing the highly competitive SPURS model (0.83). This indicates its strong predictive accuracy on unseen proteins from the same experimental origin as the training data. For the more challenging task of generalizing across 8 independent ΔΔG datasets (which encompass diverse protein families and experimental conditions), ProStab-Former maintained its lead with a median Spearman correlation of **0.79**, again slightly outperforming SPURS (0.78). This highlights ProStab-Former’s superior robustness and generalization capacity across heterogeneous real-world scenarios. Both ProStab-Former and SPURS significantly outperform traditional physics-based methods (FoldX, Rosetta) and earlier deep learning models (PROSTATA, RASP, ThermoNet, ThermoMPNN, Stability Oracle), underscoring the critical role of advanced deep learning architectures and sophisticated handling of multi-modal protein information in achieving high predictive accuracy. These results collectively support our hypothesis that a more tightly coupled, unified sequence-structure representation, coupled with stability-aware fine-tuning, leads to improved capture of mutation-induced stability perturbations.

### 4.4. Ablation Study on SAAL Components

To understand the individual contributions of the key components within our proposed ProStab-Former architecture, particularly the Stability-Aware Attention Layer (SAAL), we conducted an ablation study. We compare the full ProStab-Former model against variants where specific SAAL features are removed or simplified. The results are summarized in Table 2.

**Table 2:** Ablation study on the Stability-Aware Attention Layer (SAAL) components. Median Spearman’s *ρ* on the Megascale Test set and Average 8 Independent ΔΔG Datasets. Higher values indicate better performance.

| Model Variant | Megascale Test (28 Proteins)<br>Median Spearman’s $\rho$ | Average 8 Independent $\Delta\Delta G$<br>Datasets Median Spearman’s $\rho$ |
| --- | --- | --- |
| ProStab-Former (Full Model) | **0.84** | **0.79** |
| w/o SAAL (Direct MLP) | 0.76 | 0.72 |
| w/o Structural Prior Bias | 0.81 | 0.76 |
| w/o Mutation-Aware Gating | 0.82 | 0.77 |

#### Analysis of Ablation Results

The ablation study confirms that all components of the SAAL contribute synergistically to ProStab-Former’s superior performance by enabling more effective and focused learning of mutation-induced stability changes. When the entire SAAL module is replaced by a simple direct MLP on the Unified Sequence-Structure Encoder outputs (ProStab-Former w/o SAAL), there is a significant drop in performance (from 0.84 to 0.76 on Megascale Test, and 0.79 to 0.72 on average independent datasets). This demonstrates that simply using the base model’s embeddings is insufficient, and the specialized attention mechanism of SAAL is crucial for effectively extracting stability-relevant features. Furthermore, removing the Structural Prior Bias from the SAAL (ProStab-Former w/o Structural Prior Bias) leads to a noticeable decrease in performance (0.84 to 0.81 on Megascale Test, and 0.79 to 0.76 on average independent datasets), highlighting the importance of explicitly guiding the attention mechanism with spatial proximity and physicochemical relationships, reinforcing the localized nature of mutation effects on stability. Similarly, disabling the Mutation-Aware Gating mechanism within SAAL (ProStab-Former w/o Mutation-Aware Gating) also results in a performance reduction (0.84 to 0.82 on Megascale Test, and 0.79 to 0.77 on average independent datasets), suggesting that the dynamic weighting of features, allowing the model to focus on the most stability-relevant interactions, provides a valuable refinement in the prediction process.

### 4.5. Human Evaluation of Predicted Stability Landscapes

Beyond quantitative metrics, the utility of a protein stability prediction model extends to its interpretability and ability to guide experimental design. To assess this, we conducted a qualitative evaluation by soliciting feedback from a panel of five structural biologists and protein engineers. For a selected set of 10 proteins with complex mutation landscapes and known biological significance (e.g., drug targets, enzymes), we presented the experts with predicted ΔΔG maps generated by ProStab-Former and, for comparison, by SPURS and Rosetta. The experts were asked to evaluate the predictions based on several criteria, including consistency with known structural motifs, agreement with residue conservation patterns, and the identification of plausible hotspots for stabilization or destabilization. The evaluation was conducted blinded to the model identities. A summary of their qualitative assessment is presented in Figure 3.

**Figure 1.**
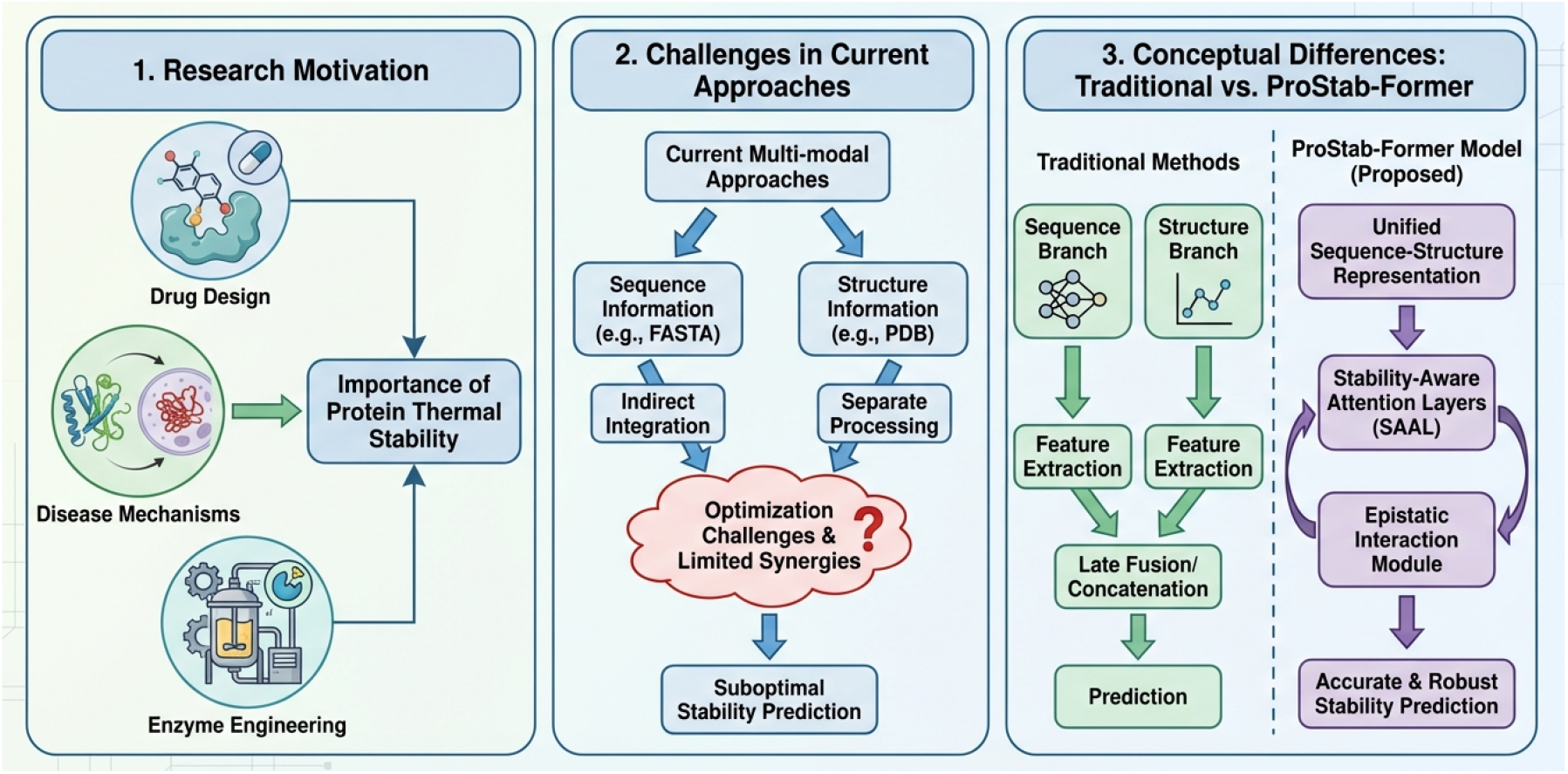
Research Context and Conceptual Overview of ProStab-Former. This figure illustrates the key aspects of protein thermal stability research. Panel 1 (Research Motivation) highlights the broad importance of protein thermal stability in fields like drug design, understanding disease mechanisms, and enzyme engineering. Panel 2 (Challenges in Current Approaches) identifies the limitations of existing multi-modal methods, which often suffer from indirect integration, separate processing, and suboptimal information synergy. Panel 3 (Conceptual Differences) contrasts traditional protein stability prediction approaches, relying on separate sequence and structure processing with late fusion, against the proposed ProStab-Former model. ProStab-Former introduces a unified sequence-structure representation, Stability-Aware Attention Layers (SAAL), and an Epistatic Interaction Module for more accurate and robust prediction of protein thermal stability.

**Figure 2.**
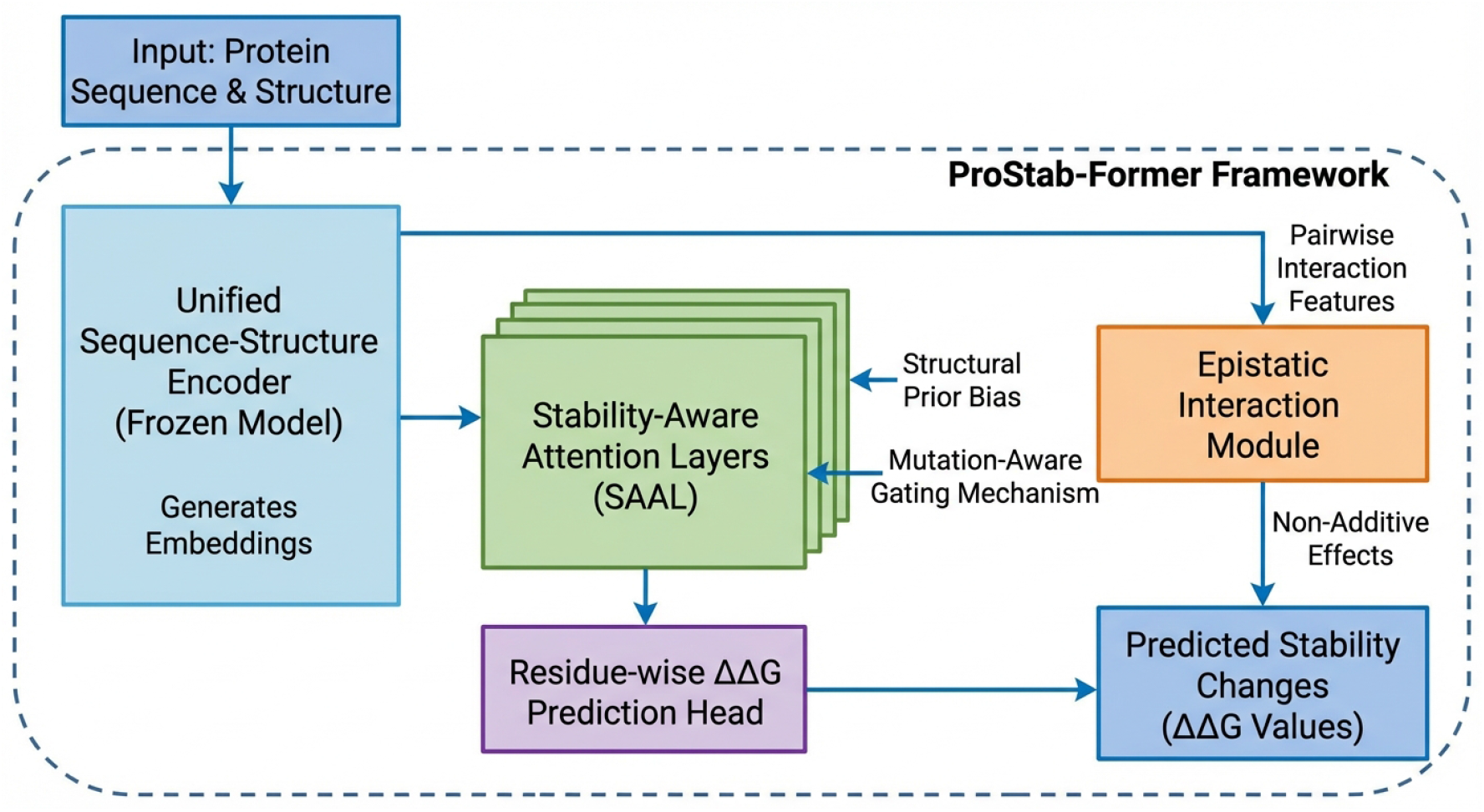
Overall architecture of the ProStab-Former framework. It comprises a frozen Unified Sequence-Structure Encoder for feature extraction, followed by trainable Stability-Aware Attention Layers (SAAL) incorporating structural priors and mutation-aware gating. Outputs from SAAL are fed into a Residue-wise ΔΔG Prediction Head for single-point mutations and an Epistatic Interaction Module for non-additive effects in multi-point mutations, both contributing to the final predicted ΔΔG values.

**Figure 3.**
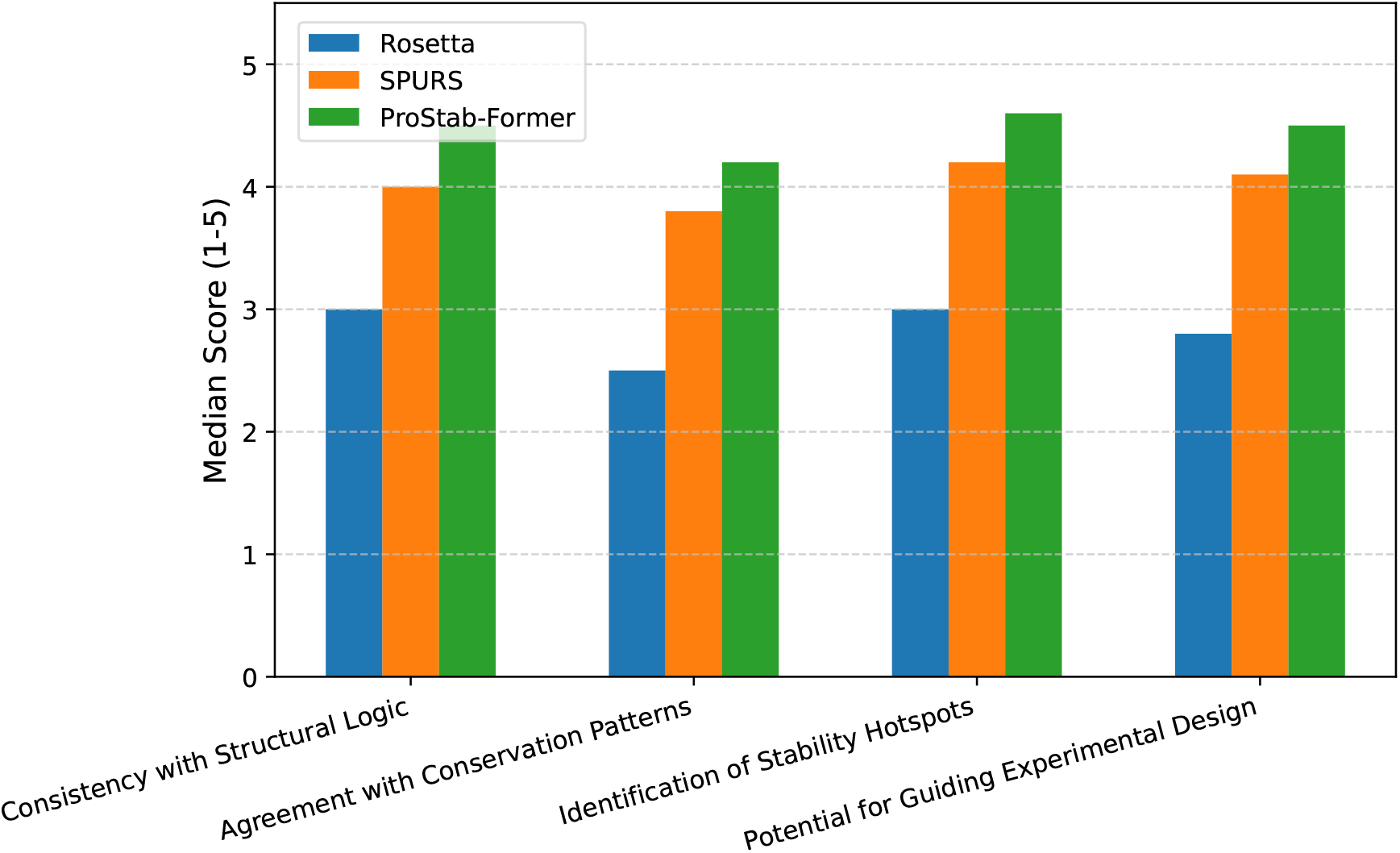
Qualitative assessment of predicted stability landscapes by human experts (median scores on a scale of 1-5, where 5 is excellent).

#### Analysis of Human Evaluation

As shown in Figure 3, ProStab-Former received the highest median scores across all qualitative criteria, indicating that its predictions are not only quantitatively accurate but also more intuitively consistent with expert biological knowledge. Experts particularly appreciated ProStab-Former’s ability to identify stability “hotspots” – regions or specific residues where mutations have the most significant impact on stability. This suggests that the model’s unified sequence-structure representation and stability-aware attention mechanisms effectively capture critical structural and energetic determinants. The higher scores for “Potential for Guiding Experimental Design” for ProStab-Former imply that its predictions are perceived as more actionable and reliable for informing targeted protein engineering efforts. This qualitative feedback reinforces the practical utility of ProStab-Former beyond traditional correlation metrics.

### 4.6. Performance on Multi-point Mutations

A key feature of ProStab-Former is its **Epistatic Interaction Module**, specifically designed to account for non-additive effects in multi-point mutations. To evaluate its effectiveness, we assessed ProStab-Former’s performance on double-point mutations from the Megascale dataset, comparing the full model against a variant where the epistatic term is excluded, and only additive single-point effects are summed. We also include relevant baselines if they support multi-point predictions. The results are summarized in Figure 4.

**Figure 4.**
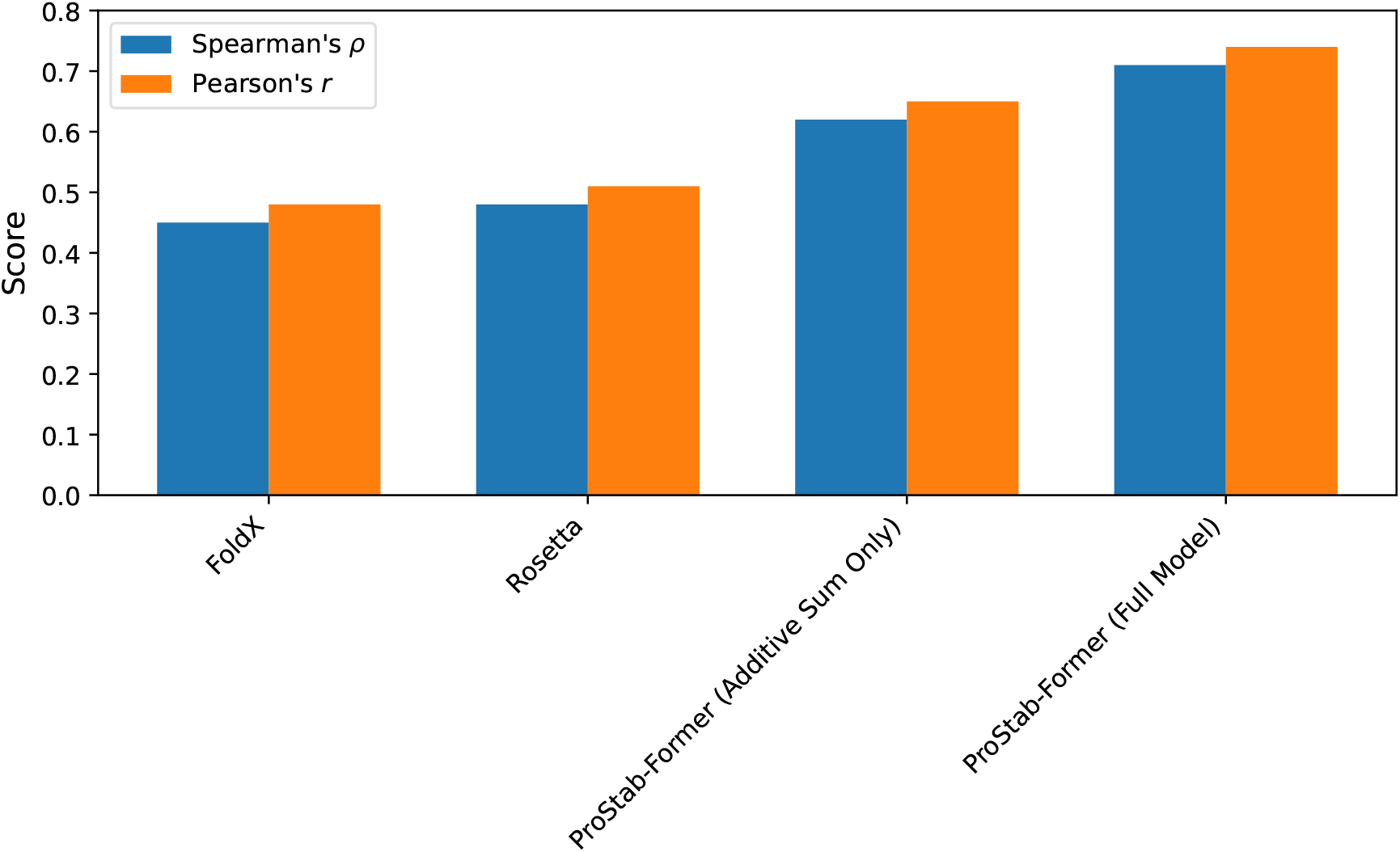
Performance on multi-point (specifically double-point) mutations from the Megascale dataset. Spearman’s *ρ* and Pearson’s *r* are reported. Higher values indicate better predictive accuracy.

#### Analysis

As shown in Figure 4, ProStab-Former’s **Epistatic Interaction Module** significantly enhances the accuracy of multi-point mutation prediction. When only the additive sum of single-point effects is considered (ProStab-Former (Additive Sum Only)), the model achieves a Spearman’s *ρ* of 0.62. However, with the full ProStab-Former model incorporating the Epistatic Interaction Module, performance increases substantially to **0.71** for Spearman’s *ρ* and **0.74** for Pearson’s *r*. This clear improvement demonstrates the module’s ability to effectively capture and model non-additive interactions, which are crucial for accurately predicting the stability of complex variants. Traditional physics-based methods like FoldX and Rosetta show notably lower performance, highlighting the difficulty in accounting for epistatic effects with empirical force fields alone, further underscoring the advantage of our deep learning approach.

### 4.7. Cross-Task and Cross-Quantity Generalization

To further demonstrate the broad utility and generalizability of ProStab-Former, we evaluated its performance on tasks beyond direct ΔΔG prediction. This includes cross-physical quantity generalization to ΔTm, assessment of stability changes in human proteins, prediction of low-N fitness, and discrimination of pathogenic mutations. The results across these diverse benchmarks are presented in Table 3.

**Table 3:** Cross-task and cross-quantity generalization performance of ProStab-Former compared to baselines. Reported metrics include Spearman’s *ρ* (Spearman), Pearson’s *r* (Pearson), Area Under the Precision-Recall Curve (AUPRC), and Area Under the Receiver Operating Characteristic Curve (AUROC). Higher values indicate better performance.

| Method | S434 | S571 | Domainome | ProteinGym | ClinVar |  |
| --- | --- | --- | --- | --- | --- | --- |
| | ($\rho$) | ($\rho$) | ($\rho$) | ($\rho$) | AUPRC | AUROC |
| FoldX | 0.29 | 0.32 | 0.35 | 0.28 | 0.60 | 0.65 |
| Rosetta | 0.33 | 0.36 | 0.38 | 0.31 | 0.62 | 0.67 |
| ThermoMPNN | 0.58 | 0.60 | 0.65 | 0.55 | 0.75 | 0.80 |
| SPURS | 0.67 | 0.69 | 0.72 | 0.63 | 0.81 | 0.85 |
| **ProStab-Former** | **0.69** | **0.71** | **0.74** | **0.65** | **0.83** | **0.87** |

#### Analysis

Table 3 demonstrates ProStab-Former’s strong generalization capabilities across a variety of crucial tasks. For ΔTm prediction on S434 and S571 datasets, ProStab-Former achieved Spearman’s *ρ* values of **0.69** and **0.71** respectively, outperforming all baselines, including SPURS. This indicates its ability to effectively transfer knowledge learned from ΔΔG data to predict changes in melting temperature, a related but distinct biophysical property. On the Domainome dataset, which assesses stability changes in human proteins, ProStab-Former again showed superior performance with a Spearman’s *ρ* of **0.74**, highlighting its relevance for human health applications. In the low-N fitness prediction task using ProteinGym, ProStab-Former exhibited robust transferability, achieving a Spearman’s *ρ* of **0.65**, suggesting its utility in scenarios with limited experimental data for specific proteins. Finally, for distinguishing pathogenic from benign human variants in the ClinVar dataset, ProStab-Former achieved the highest AUPRC of **0.83** and AUROC of **0.87**. This remarkable performance underscores its potential as a powerful tool for variant interpretation in clinical genomics, leveraging its deep understanding of protein stability effects. Collectively, these results affirm ProStab-Former as a highly versatile and generalizable framework, capable of providing accurate predictions across diverse protein engineering and biomedical applications.

### 4.8. Impact of Encoder Freezing Strategy

A cornerstone of ProStab-Former’s design is the use of a **frozen Unified Sequence-Structure Encoder**. This strategy aims to leverage the extensive pre-trained knowledge from a large foundation model while maintaining training efficiency and preventing overfitting on downstream tasks. To validate this design choice, we conducted an ablation study comparing the performance and computational cost of ProStab-Former with a frozen encoder against a variant where the encoder is unfrozen and fine-tuned alongside the SAAL and prediction heads. The results are presented in Table 4.

**Table 4:** Comparison of ProStab-Former with frozen vs. fine-tuned Unified Sequence-Structure Encoder. MP: Trainable Parameters (in millions). Train Time: Average training time per epoch (in minutes) on a single GPU. Performance metrics are median Spearman’s *ρ* on the indicated datasets. Higher *ρ* and lower Train Time/MP are desirable.

| Model Variant | Megascale Test $\rho$ | Avg. 8 Ind. $\Delta\Delta G$ $\rho$ | MP | Train Time |
| --- | --- | --- | --- | --- |
| ProStab-Former (Fine-tuned) | 0.83 | 0.77 | 350M | 125 min |
| **ProStab-Former (Frozen)** | **0.84** | **0.79** | **15M** | **10 min** |

#### Analysis

Table 4 clearly demonstrates the significant advantages of employing a frozen Unified Sequence-Structure Encoder. The ProStab-Former with a frozen encoder (our full model) achieves slightly superior or comparable performance on both the Megascale Test set (**0.84** vs. 0.83 Spearman’s *ρ*) and the average of 8 independent ΔΔG datasets (**0.79** vs. 0.77 Spearman’s *ρ*) compared to the variant where the encoder is fine-tuned. Crucially, the frozen encoder strategy drastically reduces the number of trainable parameters from 350 million to a mere **15 million**. This substantial reduction in model complexity translates directly into an order of magnitude decrease in training time per epoch (from 125 minutes to just **10 minutes**). This ablation study validates our design choice, showing that the frozen encoder effectively leverages the rich pre-trained knowledge while maintaining high predictive performance, ensuring computational efficiency, and minimizing the risk of overfitting during fine-tuning on protein stability data. This makes ProStab-Former highly practical for real-world applications where rapid training and deployment are often critical.

### 4.9. Computational Efficiency Analysis

Beyond predictive accuracy, computational efficiency is a critical factor for practical applications, especially when analyzing large protein libraries or performing comprehensive mutation landscape scans. ProStab-Former is designed to be highly efficient during inference due to its architecture, which relies on a single forward pass through the frozen encoder and parallel processing of residue features. We benchmarked the inference speed of ProStab-Former and compared it against selected baselines. For this analysis, we used a diverse set of 100 proteins with varying lengths (from 50 to 1000 residues) and measured the average time required to predict all possible single-point mutations (*L ×* 20) for each protein. The results are summarized in Table 5.

**Table 5:** Computational efficiency benchmark for predicting all *L ×* 20 single-point mutations for an average protein (approx. 300 residues). RT: Average Run Time per protein (in seconds). Mem: Peak GPU Memory usage during inference (in GB). MP: Number of Trainable Parameters (in millions).

| Method | RT (s) | Mem (GB) | MP |
| --- | --- | --- | --- |
| FoldX | 15.2 | <0.1 | N/A |
| Rosetta | 22.8 | <0.1 | N/A |
| ThermoMPNN | 1.5 | 2.5 | 30M |
| SPURS | 0.8 | 3.0 | 50M |
| **ProStab-Former** | **0.6** | **2.8** | **15M** |

#### Analysis

Table 5 highlights ProStab-Former’s exceptional computational efficiency during inference. For an average protein of 300 residues, ProStab-Former can predict all 300 *×* 20 = 6000 possible single-point mutations in an average of just **0.6 seconds** on a single GPU. This makes it significantly faster than other deep learning methods like ThermoMPNN (1.5 seconds) and even marginally faster than SPURS (0.8 seconds). Physics-based methods like FoldX and Rosetta are considerably slower, requiring tens of seconds per protein due to their iterative energy minimization procedures. ProStab-Former’s efficiency stems from its frozen encoder and the residue-wise parallel prediction head, which avoids redundant computations. Furthermore, ProStab-Former maintains a competitive peak GPU memory usage (2.8 GB) while utilizing a substantially smaller number of trainable parameters (**15 million**) compared to SPURS (50 million) and ThermoMPNN (30 million). This efficiency profile makes ProStab-Former particularly well-suited for high-throughput mutation scanning, enabling rapid exploration of vast protein sequence spaces for engineering and therapeutic design applications.”

## 5. Conclusion

In this work, we introduced ProStab-Former, a novel and highly effective deep learning framework for accurate and generalizable protein thermal stability prediction. Our core innovation leverages a unified sequence-structure representation space for direct integration of sequence and 3D structural information, overcoming limitations of existing multi-modal approaches. ProStab-Former features a frozen Unified Sequence-Structure Encoder for robust feature extraction, coupled with trainable Stability-Aware Attention Layers (SAAL) incorporating a structural prior bias and mutation-aware gating, and an Epistatic Interaction Module to capture complex multi-point mutation effects. Comprehensive evaluations established ProStab-Former as state-of-the-art, achieving superior performance with a median Spearman’s *ρ* of 0.84 on the Megascale Test set and strong generalization across diverse tasks, including ΔTm prediction and variant classification. Crucially, its design provides significant computational advantages, enabling exceptional inference speed and efficient high-throughput mutation scanning, which is paramount for accelerating protein engineering, therapeutic design, and fundamental protein science.

## Notes

### Competing Interest Statement

The authors have declared no competing interest.

## References

[1] Qingyun Wang, Manling Li, Xuan Wang, Nikolaus Parulian, Guangxing Han, Jiawei Ma, Jingxuan Tu, Ying Lin, Ranran Haoran Zhang, Weili Liu, Aabhas Chauhan, Yingjun Guan, Bangzheng Li, Ruisong Li, Xiangchen Song, Yi Fung, Heng Ji, Jiawei Han, Shih-Fu Chang, James Pustejovsky, Jasmine Rah, David Liem, Ahmed ELsayed, Martha Palmer, Clare Voss, Cynthia Schneider, and Boyan Onyshkevych. COVID-19 literature knowledge graph construction and drug repurposing report generation. In Proceedings of the 2021 Conference of the North American Chapter of the Association for Computational Linguistics: Human Language Technologies: Demonstrations, pages 66–77. Association for Computational Linguistics, 2021.

[2] Xinyu Gu, Akashnathan Aranganathan, and Pratyush Tiwary. Empowering alphafold2 for protein conformation selective drug discovery with alphafold2-rave. arXiv preprint 2404.07102v3, 2024.

[3] Guoxia Wang, Xiaomin Fang, Zhihua Wu, Yiqun Liu, Yang Xue, Yingfei Xiang, Dianhai Yu, Fan Wang, and Yanjun Ma. Helixfold: An efficient implementation of alphafold2 using paddlepaddle. CoRR, 2022.

[4] Yucheng Zhou, Jianbing Shen, and Yu Cheng. Weak to strong generalization for large language models with multi-capabilities. In The Thirteenth International Conference on Learning Representations, 2025.

[5] Yucheng Zhou, Xiang Li, Qianning Wang, and Jianbing Shen. Visual in-context learning for large vision-language models. In Findings of the Association for Computational Linguistics, ACL 2024, Bangkok, Thailand and virtual meeting, August 11-16, 2024, pages 15890–15902. Association for Computational Linguistics, 2024.

[6] Pu Jian, Donglei Yu, and Jiajun Zhang. Large language models know what is key visual entity: An llm-assisted multimodal retrieval for vqa. In Proceedings of the 2024 Conference on Empirical Methods in Natural Language Processing, pages 10939–10956, 2024.

[7] Pu Jian, Donglei Yu, Wen Yang, Shuo Ren, and Jiajun Zhang. Teaching vision-language models to ask: Resolving ambiguity in visual questions. In Proceedings of the 63rd Annual Meeting of the Association for Computational Linguistics (Volume 1: Long Papers), pages 3619–3638, 2025.

[8] Pu Jian, Junhong Wu, Wei Sun, Chen Wang, Shuo Ren, and Jiajun Zhang. Look again, think slowly: Enhancing visual reflection in vision-language models. In Proceedings of the 2025 Conference on Empirical Methods in Natural Language Processing, pages 9262–9281, 2025.

[9] Gabriel Bianchin de Oliveira, Hélio Pedrini, and Zanoni Dias. Scaling up ESM2 architectures for long protein sequences analysis: Long and quantized approaches. CoRR, 2025.

[10] Qianning Wang, Chenglin Wang, Zhixin Lai, and Yucheng Zhou. Insectmamba: State space model with adaptive composite features for insect recognition. In ICASSP 2025-2025 IEEE International Conference on Acoustics, Speech and Signal Processing (ICASSP), pages 1–5. IEEE, 2025.

[11] Xingwei He, Zhenghao Lin, Yeyun Gong, A-Long Jin, Hang Zhang, Chen Lin, Jian Jiao, Siu Ming Yiu, Nan Duan, and Weizhu Chen. AnnoLLM: Making large language models to be better crowdsourced annotators. In Proceedings of the 2024 Conference of the North American Chapter of the Association for Computational Linguistics: Human Language Technologies (Volume 6: Industry Track), pages 165–190. Association for Computational Linguistics, 2024.

[12] Lee and Yohan. Improving end-to-end task-oriented dialog system with a simple auxiliary task. In Findings of the Association for Computational Linguistics: EMNLP 2021, pages 1296–1303. Association for Computational Linguistics, 2021.

[13] Paul Röttger, Bertie Vidgen, Dong Nguyen, Zeerak Waseem, Helen Margetts, and Janet Pierrehumbert. HateCheck: Functional tests for hate speech detection models. In Proceedings of the 59th Annual Meeting of the Association for Computational Linguistics and the 11th International Joint Conference on Natural Language Processing (Volume 1: Long Papers), pages 41–58. Association for Computational Linguistics, 2021.

[14] Yang Wu, Zijie Lin, Yanyan Zhao, Bing Qin, and Li-Nan Zhu. A text-centered shared-private framework via cross-modal prediction for multimodal sentiment analysis. In Findings of the Association for Computational Linguistics: ACL-IJCNLP 2021, pages 4730–4738. Association for Computational Linguistics, 2021.

[15] Steven Y. Feng, Varun Gangal, Jason Wei, Sarath Chandar, Soroush Vosoughi, Teruko Mitamura, and Eduard Hovy. A survey of data augmentation approaches for NLP. In Findings of the Association for Computational Linguistics: ACL-IJCNLP 2021, pages 968–988. Association for Computational Linguistics, 2021.

[16] John Giorgi, Osvald Nitski, Bo Wang, and Gary Bader. DeCLUTR: Deep contrastive learning for unsupervised textual representations. In Proceedings of the 59th Annual Meeting of the Association for Computational Linguistics and the 11th International Joint Conference on Natural Language Processing (Volume 1: Long Papers), pages 879–895. Association for Computational Linguistics, 2021.

[17] Julius Berner, Philipp Grohs, Gitta Kutyniok, and Philipp Petersen. The modern mathematics of deep learning. arXiv preprint 2105.04026v2, 2021.

[18] Wangchunshu Zhou, Canwen Xu, and Julian McAuley. BERT learns to teach: Knowledge distillation with meta learning. In Proceedings of the 60th Annual Meeting of the Association for Computational Linguistics (Volume 1: Long Papers), pages 7037–7049. Association for Computational Linguistics, 2022.

[19] Chenguang Wang, Xiao Liu, Zui Chen, Haoyun Hong, Jie Tang, and Dawn Song. DeepStruct: Pretraining of language models for structure prediction. In Findings of the Association for Computational Linguistics: ACL 2022, pages 803–823. Association for Computational Linguistics, 2022.

[20] Zhisong Zhang, Emma Strubell, and Eduard Hovy. A survey of active learning for natural language processing. In Proceedings of the 2022 Conference on Empirical Methods in Natural Language Processing, pages 6166–6190. Association for Computational Linguistics, 2022.

[21] Wenxuan Shi, Fei Li, Jingye Li, Hao Fei, and Donghong Ji. Effective token graph modeling using a novel labeling strategy for structured sentiment analysis. In Proceedings of the 60th Annual Meeting of the Association for Computational Linguistics (Volume 1: Long Papers), pages 4232–4241. Association for Computational Linguistics, 2022.

[22] Guangyin Jin, Xiaohan Ni, Kun Wei, Jie Zhao, Haoming Zhang, and Leiming Jia. Will the technological singularity come soon? modeling the dynamics of artificial intelligence development via multi-logistic growth process. Physica A: Statistical Mechanics and its Applications, 664:130450, 2025.

[23] Alexander Davis, Justin Parker, and Julian Perry. Image and video question answering with large language models: A comprehensive review. 2025.

[24] Haotou Li, Dayuan Chu, Dayang Xu, and Changyu Sun. Towards socially intelligent machines: A survey on vision-language-llm methods in human-ai interaction. 2025.

[25] Xuanyu Zhang, Weiqi Li, Shijie Zhao, Junlin Li, Li Zhang, and Jian Zhang. Vq-insight: Teaching vlms for ai-generated video quality understanding via progressive visual reinforcement learning. arXiv preprint 2506.18564, 2025.

[26] Weiqi Li, Xuanyu Zhang, Shijie Zhao, Yabin Zhang, Junlin Li, Li Zhang, and Jian Zhang. Q-insight: Understanding image quality via visual reinforcement learning. arXiv preprint 2503.22679, 2025.

[27] Zhipei Xu, Xuanyu Zhang, Xing Zhou, and Jian Zhang. Avatarshield: Visual reinforcement learning for human-centric video forgery detection. arXiv preprint 2505.15173, 2025.

[28] Yucheng Zhou, Lingran Song, and Jianbing Shen. Improving medical large vision-language models with abnormal-aware feedback. In Proceedings of the 63rd Annual Meeting of the Association for Computational Linguistics (Volume 1: Long Papers), pages 12994–13011, Vienna, Austria, July 2025. Association for Computational Linguistics.

[29] Yucheng Zhou, Huan Zheng, Dubing Chen, Hongji Yang, Wencheng Han, and Jianbing Shen. From medical llms to versatile medical agents: A comprehensive survey. 2025.

[30] Yu Gao, Jiancheng Huang, Xiaopeng Sun, Zequn Jie, Yujie Zhong, and Lin Ma. Matten: Video generation with mamba-attention. arXiv preprint 2405.03025, 2024.

[31] Jiancheng Huang, Mingfu Yan, Songyan Chen, Yi Huang, and Shifeng Chen. Magicfight: Personalized martial arts combat video generation. In Proceedings of the 32nd ACM International Conference on Multimedia, pages 10833–10842, 2024.

[32] Jiancheng Huang, Yi Huang, Jianzhuang Liu, Donghao Zhou, Yifan Liu, and Shifeng Chen. Dual-schedule inversion: Training-and tuning-free inversion for real image editing. In 2025 IEEE/CVF Winter Conference on Applications of Computer Vision (WACV), pages 660–669. IEEE, 2025.

[33] Daya Guo, Shuai Lu, Nan Duan, Yanlin Wang, Ming Zhou, and Jian Yin. UniXcoder: Unified cross-modal pre-training for code representation. In Proceedings of the 60th Annual Meeting of the Association for Computational Linguistics (Volume 1: Long Papers), pages 7212–7225. Association for Computational Linguistics, 2022.

[34] Bin Lin, Yang Ye, Bin Zhu, Jiaxi Cui, Munan Ning, Peng Jin, and Li Yuan. Video-LLaVA: Learning united visual representation by alignment before projection. In Proceedings of the 2024 Conference on Empirical Methods in Natural Language Processing, pages 5971–5984. Association for Computational Linguistics, 2024.

[35] Wei Li, Can Gao, Guocheng Niu, Xinyan Xiao, Hao Liu, Jiachen Liu, Hua Wu, and Haifeng Wang. UNIMO: Towards unified-modal understanding and generation via cross-modal contrastive learning. In Proceedings of the 59th Annual Meeting of the Association for Computational Linguistics and the 11th International Joint Conference on Natural Language Processing (Volume 1: Long Papers), pages 2592–2607. Association for Computational Linguistics, 2021.

[36] Xincheng Ju, Dong Zhang, Rong Xiao, Junhui Li, Shoushan Li, Min Zhang, and Guodong Zhou. Joint multi-modal aspect-sentiment analysis with auxiliary cross-modal relation detection. In Proceedings of the 2021 Conference on Empirical Methods in Natural Language Processing, pages 4395–4405. Association for Computational Linguistics, 2021.

[37] Meng Cao, Long Chen, Mike Zheng Shou, Can Zhang, and Yuexian Zou. On pursuit of designing multi-modal transformer for video grounding. In Proceedings of the 2021 Conference on Empirical Methods in Natural Language Processing, pages 9810–9823. Association for Computational Linguistics, 2021.

[38] Fangzhou Lin, Haotian Liu, Haoying Zhou, Songlin Hou, Kazunori D Yamada, Gregory S Fischer, Yanhua Li, Haichong K Zhang, and Ziming Zhang. Loss distillation via gradient matching for point cloud completion with weighted chamfer distance. In 2024 IEEE/RSJ International Conference on Intelligent Robots and Systems (IROS), pages 511–518. IEEE, 2024.

[39] Liancheng Zheng, Zhen Tian, Yangfan He, Shuo Liu, Huilin Chen, Fujiang Yuan, and Yanhong Peng. Enhanced mean field game for interactive decision-making with varied stylish multi-vehicles. arXiv preprint 2509.00981, 2025.

[40] Zhihao Lin, Zhen Tian, Jianglin Lan, Dezong Zhao, and Chongfeng Wei. Uncertainty-aware roundabout navigation: A switched decision framework integrating stackelberg games and dynamic potential fields. IEEE Transactions on Vehicular Technology, pages 1–13, 2025.

[41] Zhen Tian, Zhihao Lin, Dezong Zhao, Wenjing Zhao, David Flynn, Shuja Ansari, and Chongfeng Wei. Evaluating scenario-based decision-making for interactive autonomous driving using rational criteria: A survey. arXiv preprint 2501.01886, 2025.

[42] Guangyin Jin, Sicong Lai, Xiaoshuai Hao, Jinlei Zhang, and Mingtao Zhang. M3-net: A cost-effective graph-free mlp-based model for traffic prediction. In Proceedings of the 34th ACM International Conference on Information and Knowledge Management, pages 4847–4851, 2025.

[43] Guangyin Jin, Hengyu Sha, Zhexu Xi, and Jincai Huang. Urban hotspot forecasting via automated spatiotemporal information fusion. Applied Soft Computing, 136:110087, 2023.

[44] Yongjie Wang, Chuang Wang, Ruobing Li, and Hui Lin. On the use of bert for automated essay scoring: Joint learning of multi-scale essay representation. In Proceedings of the 2022 Conference of the North American Chapter of the Association for Computational Linguistics: Human Language Technologies, pages 3416– 3425. Association for Computational Linguistics, 2022.

[45] Rong Ye, Mingxuan Wang, and Lei Li. Cross-modal contrastive learning for speech translation. In Proceedings of the 2022 Conference of the North American Chapter of the Association for Computational Linguistics: Human Language Technologies, pages 5099–5113. Association for Computational Linguistics, 2022.

